# Melaninization Reduces *Cryptococcus neoformans* Susceptibility to Mechanical Stress

**DOI:** 10.1101/2022.09.01.506291

**Authors:** Ellie Rose Mattoon, Radames JB Cordero, Arturo Casadevall

## Abstract

Melanin is a complex pigment found in various fungal species that is associated with a multitude of protective functions against environmental stresses. In *Cryptococcus neoformans*, melanin is synthesized from exogenous substrate and deposited in the cell wall. Although melanin is often cited as a protector against mechanical stress, there is a paucity of direct experimental data supporting this claim. To probe whether melanin enhances cellular strength, we used ultrasonic cavitation and French pressure cell press to stress cryptococcal cells and then measured changes in cellular morphology and survival for melanized and non-melanized *C. neoformans*. Melanized yeast exhibited lower rates of fragmentation and lower decreases in cell area when compared to non-melanized yeast after sonication and French press conditions. Our results indicate that melanization protects against some of the morphologic changes initiated by mechanical energy derived from either sonic cavitation or French press, thus supporting the notion that this pigment provides mechanical strength to fungal cell walls.

**Importance:** Melanin has been shown from prior experiments in microbiology to be associated with protection against environmental stressors and has often been cited as being associated with mechanical stress protection. However, there is a lack of direct experimentation to confirm this claim. By examining the response of melanized and non-melanized *C. neoformans* to sonication and French press, we report differences in outcomes dependent not only based on melanization status but also culture age. Such findings have important implications in the design and interpretation of laboratory experiments involving *C. neoformans*. In addition, uncovering some of melanin’s mechanical properties promotes further research into fungal melanin’s applications in healthcare and industry.

## Introduction

Melanin is a complex multifunctional pigment found in the cell wall and extracellular vesicles of many fungal species. This pigment is associated with a multitude of protective functions against stresses such as ultraviolet radiation, desiccation, and toxic heavy metals (1). Although melanin has often been cited as a protector against mechanical stress (1), there is scant data to directly support this claim. It is known that melanotic fungi can survive in environments at extremely low and extremely high pressures (extraterrestrial conditions and deep-sea hydrothermal vents, respectively (2–4). In addition, melanin contributes to virulence, and fungal pathogens likely undergo compressive stresses when invading host tissues (5,6). The invasive hyphal growth of *Wangiella dermatitidis* in agar exhibited dependence on melanin synthesis (7), and melanin is associated with higher turgor pressures and increased cell wall rigidity in *Gaeumannomyces graminisvar*.*graminis* hyphopodia (8). In addition, melanized *C. neoformans* cells were more likely to survive orbital flight and microgravity conditions than non-melanized cells (9). Melanization prevents liposome transit in *C. neoformans*, a process that is also consistent with increased rigidity (10). Lastly, melanin has been associated with osmotic stress resistance in black yeast (11), which other studies have linked to properties of the fungal cell wall which allow cells to rapidly expand in hypoosmotic environments to and avoid rupture (12).

Sonication is a physical process that fragments cellular samples using ultrasonic vibrations, which create cavitating bubbles in liquid that release elastic waves and eddies as they collapse (13). As eddies interact with cells, they create a pressure difference, which can cause the cell wall to disintegrate (13). At lower power settings and shorter durations, sonication is often used to remove cell clumps from a sample with minimal effects on viability (14). However, an assessment of *Saccharomyces cerevisiae* exposed to sonication posited a fungicidal effect on yeast cells in close proximity to cavitation bubbles, while other cells exhibited a fungistatic effect that increased with longer durations of sonication (15). Earlier assessments of sonic energy’s effects on *Saccharomyces ellipsoideus* noted that the majority of cells viewed under the microscope were fragmented (16). Recently our group used sonication to remove the *Cryptococcus neoformans* polysaccharide capsule (17) but did not probe the effects on the cell wall.

The French pressure cell press, often referred to as simply the French press, is a device used to disrupt the membrane of cells, often for purposes of protein extraction (18). Unlike sonication stress, cells passed through a French press rupture when being passed through a narrow valve at high pressures (19). Cellular organelles often remain intact during this process (18). In addition, cells passed through a French press are subjected to a single instance of maximum shear force, while sonicated samples may be subject to multiple mechanical shockwaves (18). Like sonication, we have used the French press to strip the capsule from *C. neoformans* (17).

While melanized fungal cells are relatively more resistant to chemical and electromagnetic radiation stress, the evidence that it contributes to cellular mechanical strength is indirect (1). Hence, we sought to obtain direct evidence for this widely held supposition. In this study, we investigated the response of melanized and non-melanized *Cryptococcus neoformans* to sonication and French press stress with the goal of determining if outcomes differed between the two groups. Ultimately, melanized cells were more resistant than non-melanized cells to ultrasonic cavitation and mechanical shear stresses.

## Results

### Cellular Mass Decreases Post-Sonication

Previous researchers have made the observation that the cellular pellet of sonicated samples are visually smaller than the cellular pellet of unsonicated controls, a finding attributed to stripping of the polysaccharide capsule (20). We also observed this phenomenon during experimentation in both melanized and non-melanized H99 (**Figure 1d-e**). To ascertain whether this visual observation was due to a change in volume or mass, we washed samples and controls in microcentrifuge tubes and recorded their mass. For both melanized and non-melanized samples, we observed a significant decrease in mass when comparing sonicated and unsonicated samples (**Figure 1a**). Given prior results that the capsule accounts for up to 90% of *C. neoformans* cell volume (21) and that the process of sonication stripped this capsule (17), we repeated the experiment with cap59 acapsular mutants. We hypothesized that these would not undergo change, and no significant different in cellular mass was obtained (**Figure 1b**). In addition, the mass of the sonicated H99 culture was similar to the mass of all cap59 populations. As these mutants lack capsules, the absence of a mass difference pre- and post-sonication is consistent with the mass decrease in H99 arising from removal of its cellular capsule. The remainder of the sonicated and control cultures were plated; while colonies grew on the control plates, no colonies grew on the sonicated plates (data not shown), consistent with a fungicidal effect from our sonication settings. When this experiment was repeated with a lethal heat shock; there was no significant difference in cellular mass (**Figure 1c**) or capsule radius (**Figure 1d**), which suggests that cellular death from our sonication treatment was not a major contributor to the loss of the polysaccharide capsule or a decrease in mass.

**Figure 1:**
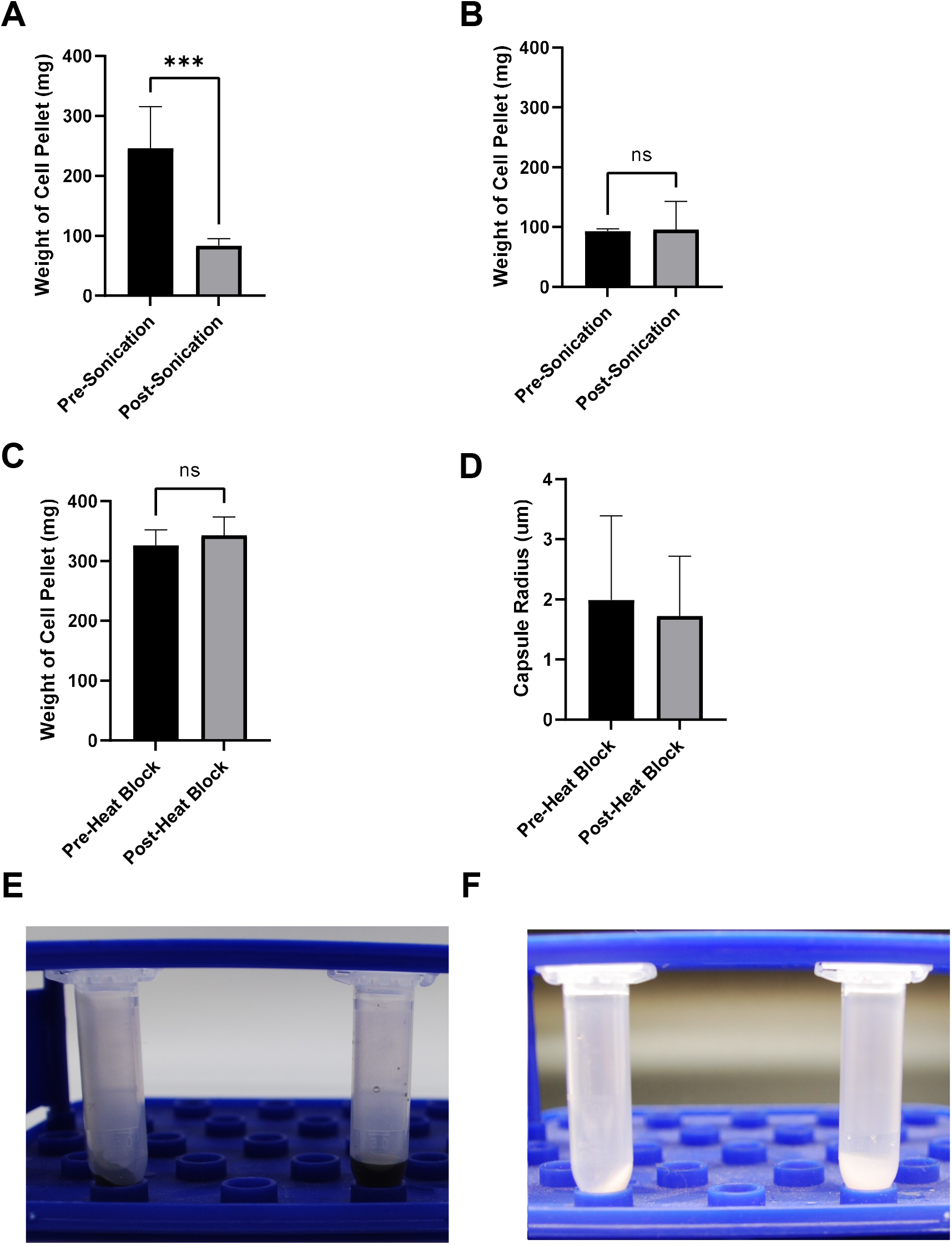
Change in Cellular Mass Post-Sonication. (A) Average mass of twice-washed cell pellets of melanized and non-melanized H99 with and without sonication. Cell pellets contained approximately 2 ×10^8^ cells. (B) Average mass of twice-washed cell pellets of melanized and non-melanized cap59 with and without sonication. Cell pellets contained approximately 2 ×10^8^ cells. (C) Average mass of twice-washed cell pellets of melanized and non-melanized H99 before and after being placed on a heat block at 75 °C for 30 minutes. Cell pellets contained approximately 1.5 × 10^8^ cells. (D) Average capsule radius of melanized and non-melanized H99 before and after being placed on a heat block at 75°C for 30 minutes. (E) Side-by-side comparison of melanized H99 sonicated (left) or unsonicated (right) cell pellets. (F) Side-by-side comparison of non-melanized H99 sonicated (left) or unsonicated (right). Experiments were repeated twice with comparable results. In all panels *, **, ***, **** represent P < 0.05, 0.01, 0.001, and .0001 respectively.

### Non-melanized Cells Are More Likely to Rupture Due to Sonication

To examine the effects of melanization on cellular susceptibility to fragmentation under sonication stress, melanized and non-melanized cells that had been sonicated were examined by light microscopy. At the specified amplitude, the culture temperature did not rise above 40 °C, likely not impacting viability by heat stress (data not shown). Fragmentation was defined and counted in all samples (**Figure 2**). For each sample, we counted 200 cells and recorded the proportion of cells that met the criteria for fragmentation (**Figure 2**). When considering results with all experiments and with at all culture ages, non-melanized H99 and cap59 cells exhibited significantly greater proportions of fragmentation under the microscope (**Figure 3a-b**). To examine the effects of sonication on cell area, brightfield microscopy images of H99 stained with methylene blue were analyzed in ImageJ. Magnification or field of view was not adjusted when examining cultures of the same age to ensure comparable area comparisons. Melanized cells exhibited higher cell areas than non-melanized cells post-sonication (**Figure 4a-c**). However, at all culture ages, control melanized and non-melanized cells manifested non-significant differences in cell area (p=0.8223, 0.3518, and 0.8152, respectively).

**Figure 2:**
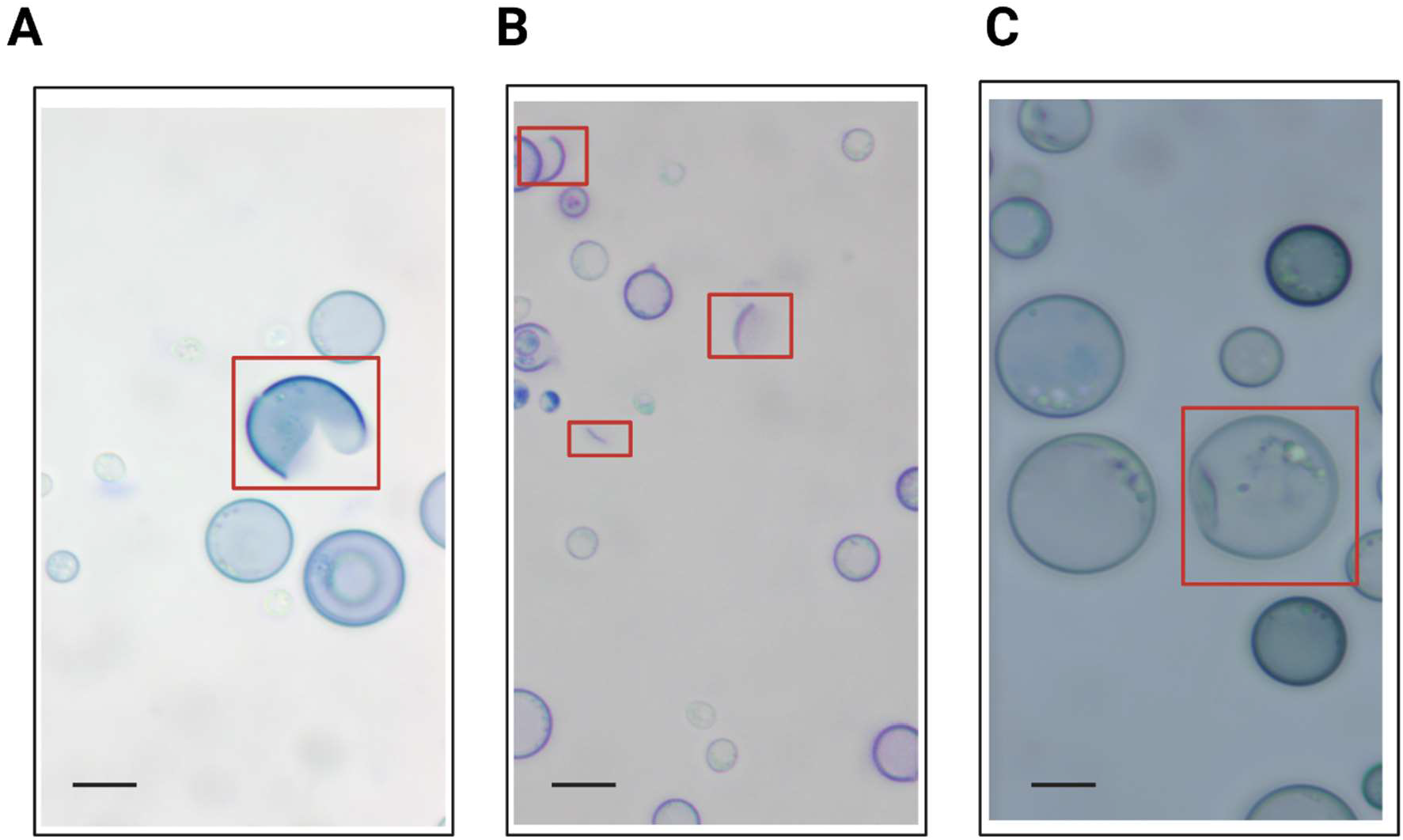
Cellular Ruptures Defined Using Brightfield Microscopy. (A) “Pacman” cells were mostly intact except for a portion of the cell wall that appeared to be missing. (B) Crescent-shaped fragments appeared as stained arcs with a circumference less than half of that of an intact cell. (C) Deformed or “wrinkled peas” cells were identified in both control and treatment cultures and as a result were excluded from analysis. Microscopic images were taken using brightfield microscopy at 100x magnification and methylene blue staining. All images shown are of five-day cell cultures that underwent sonication. Comparable morphologies were identified in cells that had undergone French press. Scale Bar 6 um.

**Figure 3:**
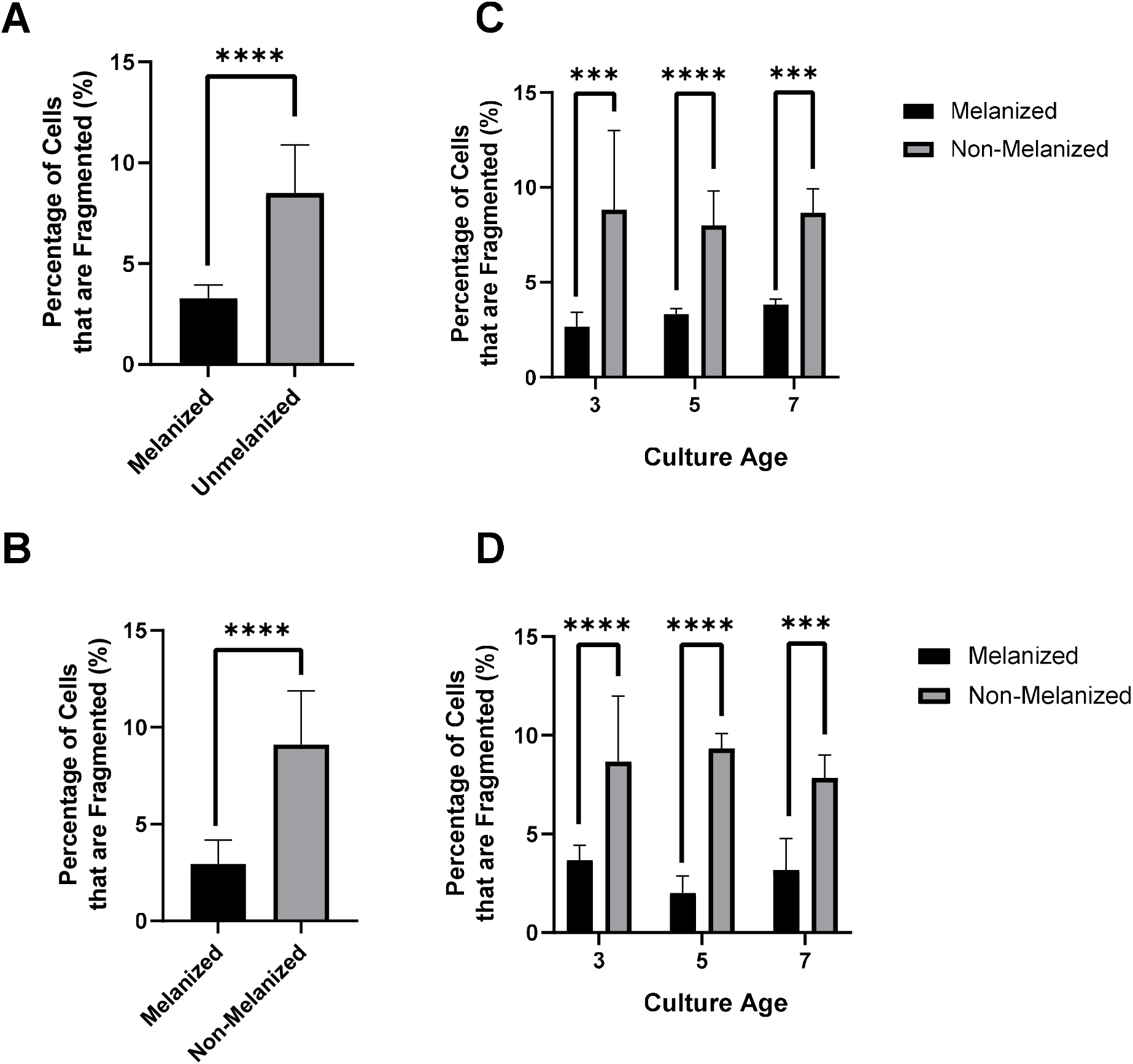
Cell Fragmentation Following Sonication. (A) Percent fragmentation of melanized and non-melanized H99 that underwent three 30-second sonication pulses (n=1800 for each column) (B) Percent fragmentation of melanized and non-melanized cap59 that underwent three 30-second sonication pulses (n=1800 for each column) (C) Percent fragmentation of melanized and non-melanized H99 that underwent three 30-second sonication pulses three days, five days, and seven-days post-inoculation (n=600 for each column). (D) Percent fragmentation of melanized and non-melanized cap59 that underwent three 30-second sonication pulses three days, five days, and seven days post-inoculation (n=600 for each column). Experiments were repeated twice for cap59 and three times for H99 with comparable results. In all panels, *, **, ***, **** represent P < 0.05, 0.01, 0.001, and .0001 respectively.

**Figure 4:**
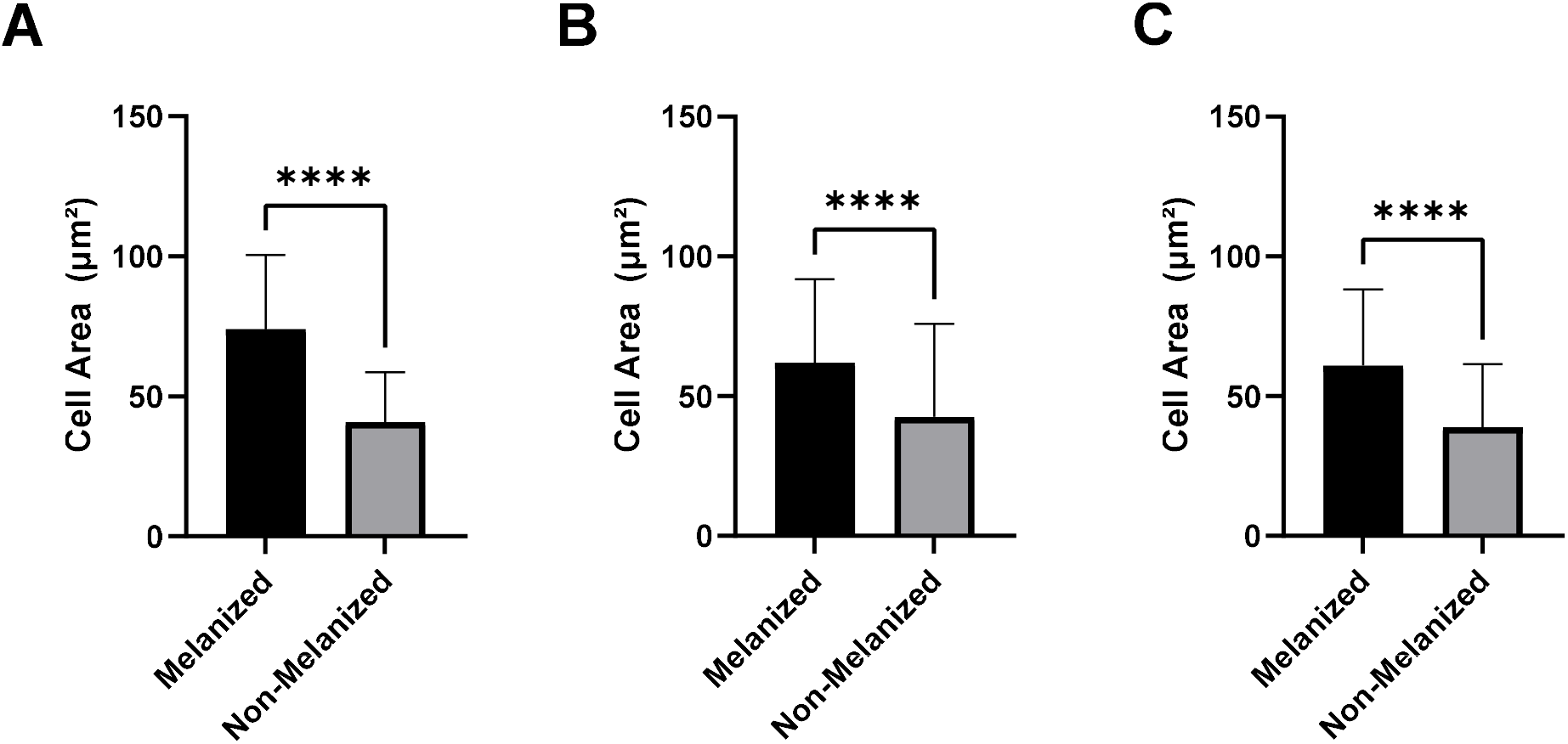
Cell Area Changes from Sonication Experiments. (A) Cell area of both melanized and non-melanized H99 three days post-inoculation that underwent three 30-second sonication pulses (n=100 for each column). (B) Cell area of both melanized and non-melanized H99 five days post-inoculation that underwent sonication (n=100 for each column). (C) Cell area of both melanized and non-melanized H99 seven days post-inoculation that underwent sonication (n=100 for each column). In all panels, *, **, ***, **** represent P < 0.05, 0.01, 0.001, and .0001 respectively.

### Non-melanized Cells are More Likely to Rupture After French Press

Both melanized and non-melanized cells passed through a French press were examined under the microscope. Fragmentation was defined and counted as described in **Materials and Methods** (**Figure 2**). For cells of all culture ages, non-melanized cells also exhibited significantly greater proportions of fragmentation under the microscope (**Figure 5a**). When stratifying the results by culture ages, the statistical magnitude of the difference increased with increasing culture age (**Figure 5b**). An increase in fragmentation over time was observed in all cultures that had undergone French Press, regardless of melanization status (**Figure 5c**). Melanized cells exhibited higher cell areas than non-melanized cells post-French press (**Figure 6a-c**); this significance increased with culture age (**Figure 6a-c**). However, at all culture ages, melanized and non-melanized cells manifested non-significant differences in cell area (p=0.2388, 0.2152, and 0.9344, respectively).

**Figure 5:**
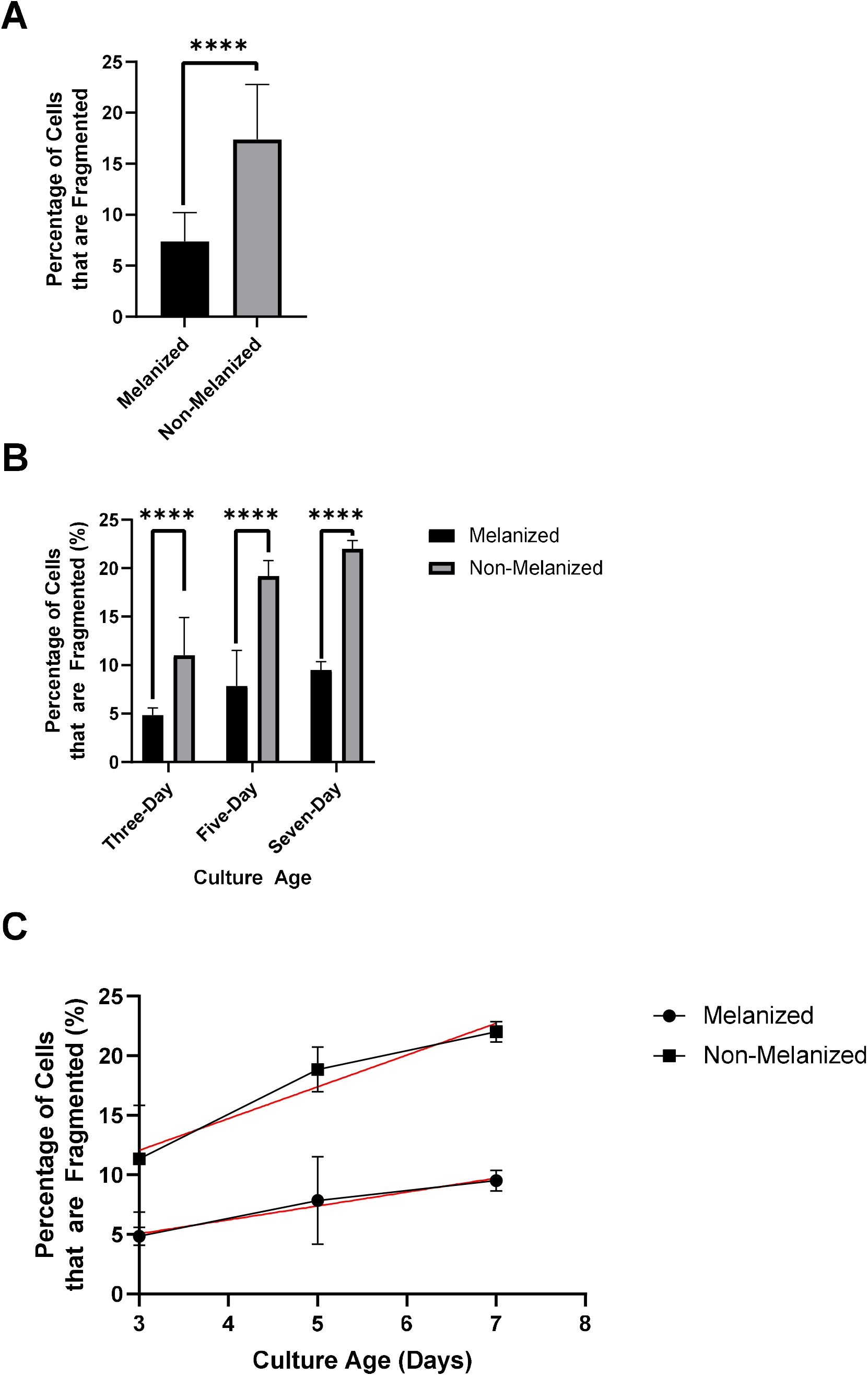
Cell Fragmentation Following French Press. (A) Percent fragmentation of melanized and non-melanized H99 that underwent French press at 4964 kPa (n=1800 for each column). (B) Percent fragmentation of melanized and non-melanized H99 that underwent French Press at 4964 kPa three days, five days, and seven days post-inoculation (n=600 for each column). Experiments were repeated three times with comparable results. (C) Percent fragmentation of melanized and non-melanized H99 that underwent French Press at 4964 kPa three, five-, and seven-days post-inoculation (n=600 for each data point), regrouped to display trends with respect to culture age. Trends were determined via linear regression. For melanized cultures, slope=1.167. For non-melanized cultures, slope=2.667. In all panels *, **, ***, **** represent P < 0.05, 0.01, 0.001, and .0001 respectively.

**Figure 6:**
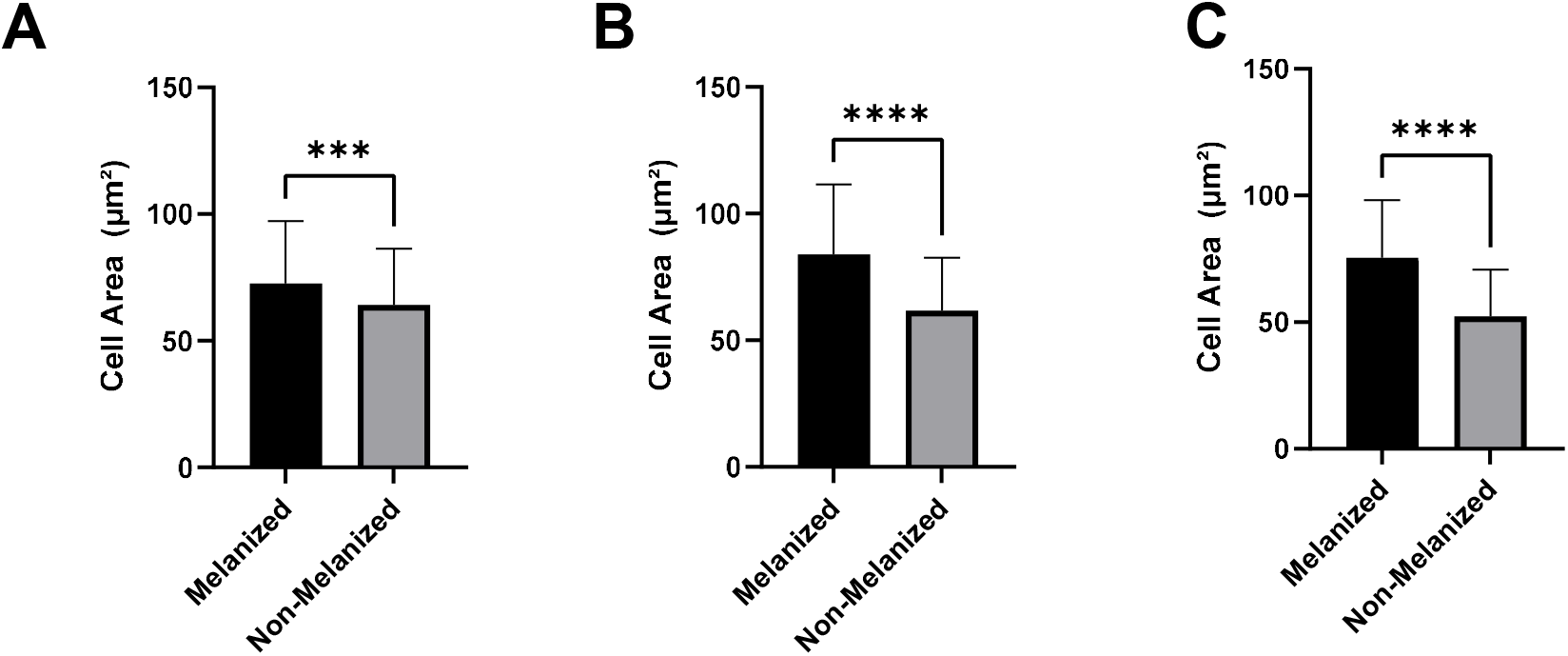
Cell Area after French Press procedure. (A) Cell area of both melanized and non-melanized H99 three days post-inoculation that underwent French Press at 4964 kPa (n=200 for each column). (B) Cell area of both melanized and non-melanized H99 five days post-inoculation that underwent French Press at 4964 kPa (n=200 for each column). (C) Cell area of both melanized and non-melanized H99 seven days post-inoculation that underwent French Press at 4964 kPa (n=200 for each column); *, **, ***, **** represent P < 0.05, 0.01, 0.001, and .0001 respectively.

## Discussion

In this study, we examined the widely believed assumption that melanization protected fungal cells from mechanical stress using melanized and non-melanized C. *neoformans* yeast and obtained evidence confirming its veracity. Melanin has been shown in prior experiments to have a protective effect against radiative (22–24), oxidative (25), temperature (26–28), and some forms of mechanical stress (7,8). Determination of melanin’s protective properties during sonication and French press induced shear stress holds implications for further understanding of melanin’s role in the *C. neoformans* response to mechanical stressors in laboratory experiments.

Exposure to ultrasound waves caused changes to the cellular mass and morphology of H99; likewise, French press caused changes to H99 cell area and morphology. While changes to cellular mass and/or area appeared to be constant regardless of melanization status, melanized cells were less likely to fragment in response to sonication. While fragmentation is a sufficient cause of cell death from sonication, this is not necessary for cellular death to occur. A prior study found that melanized cells survived less than non-melanized cells at lower sonication amplitudes (20), suggesting that while melanin may protect against physical disruption, its protective effects may not apply to cavitation’s fungicidal effects on intact cells.

Melanin is not generally recovered from *C. neoformans* cells until the cultures reach stationary phase after about three days of growth, and cells continue to accumulate melanin as late as eight days post-inoculation (27). Our observations demonstrating that melanization increases in its protective effect with culture age following French press is consistent with the notion that this phenomenon is correlated with larger qualities of melanin in culture. Sonication can be used to remove the polysaccharide capsule of *C. neoformans* (17), and the supernatant of washed sonicated cultures has been shown to have higher carbohydrate concentrations than control cultures (20). These results suggest that most of the mass loss during sonication was from the removal of the capsule. However, we also report that, regardless of melanization status, *C. neoformans* generally becomes more susceptible to mechanical stresses with increasing culture age. We hypothesized that cells would become more brittle as they aged, possibly due to the changing composition of the cell wall (29). In addition, these findings are consistent with prior studies that culture age reduces resistance to stress. For example, non-melanized *C. neoformans* had lower survival rates against UV radiation at higher culture ages (2,22). One previous study that noted decreasing thermotolerance in older melanized cultures attributed the phenomenon to a decrease in cell wall fluidity due to the continuous deposition of melanin over time (27). Other studies have noted that prolonged growth in stationary phase is associated with increased α-1,3-glucan expression in the capsule and changing elasticity in *C. neoformans* (30). The notion that *C. neoformans* experiences changes in its mechanical properties as culture ages has important implications in experiment design.

Although one must always be cautious in making mechanistic inferences across large size scales, we note that our microscopic observations that melaninization confers structural strength onto melanized cells relative to non-melanized cells is consistent with what we know about the effects of cell wall melanin deposition in *C. neoformans*. Melanization is known to lead to the formation of covalent bonds between the pigment and cell wall polysaccharide components (31) and chitosan (32), suggesting the formation of cross-linked lattices that could confer increased structural strength to the cell wall. Dopamine-mediated scleratinization of chitin resulting in the formation of melanin-like material was associated with a 2.5-fold increase in tensile strength (33). Furthermore, progressive melanization reduces the pore size of *C. neoformans* cell wall (34) and the filling of such spaces could provide additional structural strength.

In summary, melanized cells were less vulnerable to disruption by sonication and French press than non-melanized cells. Melanin is known to be a chemically tough polymer that resists acid degradation and its presence in cell wall could provide tensile strength that would reduce the vulnerability of melanized cells to sonication and French press relative to non-melanized cells. Our results provide direct evidence for the notion that melanin gives cells greater mechanical stability when placed in cell walls.

## Materials and Methods

### Species, strains, and media used in this work

*C. neoformans* strain H99 and cap59 were kept frozen in 15% glycerol stocks and precultured onto Sabouraud dextrose broth for 48 hours prior to inoculation in minimal media at 30 °C. Minimal media contained 15 mM dextrose, 10 mM MgSO4, 29.3 mM KH2PO4, 13 mM glycine, 3 mM thiamine-HCL; adjusted to pH 5.5) with or without 1 mM of L-Dopa to induce melanization.

### Sonication

Melanized and non-melanized cultures were analyzed at 3, 5, and 7 d. The yeast cells were counted with a hemocytometer, washed twice with minimal media, adjusted to 10^8^ cells/ml and added to glass cuvettes in volumes of 3 mL. Cuvettes were sonicated in a Horn sonicator (Fisher Scientific Sonic Dismembrator Model 100, Waltham, MA) at a power of 22 Watts over an ice bath for three 30-second pulses. Every sonication experiment conducted featured two control groups. One control group remained in an ice bath for the duration of the sonication process. Another control group remained at room temperature.

### French Press

Melanized and non-melanized cultures were analyzed at 3, 5, and 7 d of age. The yeast cells were counted with a hemocytometer, washed twice with minimal media, adjusted to 10^7^ cells/ml and added to Falcon graduated tubes in volumes of 20 ml. Samples were placed through the French press G-M High Pressure Standard Cell (Clifton, NJ) once at a setting of 4964 kilopascal (kPa) or 720 pounds per square inch (psi). One melanized control and one non-melanized control was not placed through the French press.

### Cellular Changes in Mass

After sonication at an amplitude of eight, 1.0 mL each of treatment and control cultures were washed with 1 mL minimal media in 2 mL microcentrifuge tubes. The supernatant was removed, and the tubes were weighed on a scale. Five empty microcentrifuge tubes were weighed, and the average of these weights was used to determine the final weight of cell pellets.

To analyze the effect of heat death on cellular mass, cells were killed on a heat block at 75 °C for 35 minutes. Following heat death, treatment and control cultures were washed with 1 mL minimal media in 2 mL microcentrifuge tubes. The supernatant was removed, and the tubes were weighed on a scale. Heat-killed cells were also mixed with india ink and imaged on an Olympus AX70 microscope using QImaging Retiga 1300 digital camera and the QCapture Suite V2.46 software (QImaging). Capsule measurements were obtained using a Quantitative Capture Analysis program developed by the lab (35).

### Analysis of Cellular Rupture

After sonication or French Press, cultures were directly added to slides with methylene blue and imaged with an Olympus AX70 microscope using QImaging Retiga 1300 digital camera and the QCapture Suite V2.46 software (QImaging). To visually define cellular fragments, multiple brightfield microscopy images of 5-day cultures were randomly taken at 100x magnification. To quantify cellular fragments, brightfield microscopy images were taken of all four corners and the center of the slide at 20x magnification. The number of cellular fragments for 200 cells was then determined visually, starting with the top left corner of the slide and then moving to the top right, bottom right, bottom left, and center if more cells were needed. This was completed in triplicate for both melanized and non-melanized cultures, leading to a total of 600 cells counted per treatment condition. Control cultures and, in the case of the sonication experiment room temperature cultures, were examined to confirm the absence of cellular fragments.

### Analysis of Cellular Area

Cell area analyses were completed using methods previously described (36) in ImageJ (37).

### Data Analysis

Data analysis was completed in *Prism* (38).

